# Distinct DNA metabolism and anti-proliferative effects of goat urine metabolites: An explanation for xeno-tumor heterogeneity

**DOI:** 10.1101/815548

**Authors:** Ajay Kumar, Swati Swami, Nilesh Kumar Sharma

**Author notes:** Corresponding author: Dr. Nilesh Kumar Sharma Associate Professor, Cancer and Translational Research Lab Department of Biotechnology, Dr. D. Y. Patil Biotechnology & Bioinformatics Institute, Pune Dr. D. Y Patil Vidyapeeth Pune, Pune, MH, 411033.

## Abstract

**Background:** The tumor microenvironment, including microbiome populations in the local niche of several types of solid tumors like mammary and colorectal cancer are distinct. The occurrence of one type of cancer over another varies from animals to human individuals. Further, clinical data suggest that specific cancer types such as mammary and colorectal cancer are rare in ruminant like goat.

**Methods:** Fresh urine samples were collected from healthy ruminates including cow, goat, buffalo, ox, horse, jenny and human and subjected to fractionation using drying, vortexing, centrifugation and sterile filtration in DMSO solvent. Collected urine DMSO fraction (UDF) samples from all sources were subjected DNA metabolizing assay with plasmid DNA pBR322 and genomic DNA of MCF-7 cells. Further, based on the discernible DNA metabolizing effects, goat UDF was tested for anti-proliferative effects upon HCT-116 and MCF-7 cells using Trypan blue due exclusion assay.

**Results:** This paper reports that goat UDF possesses very clear DNA metabolizing effects (up to 95%) upon plasmid and genomic DNA compared to other ruminants and human UDF samples. Interestingly autoclaving of goat UDF and other sample results in the significant loss of DNA metabolizing effects. In this way, data potentially indicate that the goat UDF sample contains metabolite or similar organic compounds. Further, in vitro treatment of the goat UDF sample shows discernible anti-proliferative effects upon HCT-116 (up to 75%) and MCF-7 (up to 40%).

**Conclusion:** This study signifies the clear differences in DNA metabolizing effects of goat UDF and well correlated with anti-proliferative effects upon HCT-116 and MCF-7 cells. This study is of first report to show the comparison of urine metabolites and an indirect link to support the possible reasons behind xeno-tumor heterogeneity as rare occurrences of colorectal and mammary cancer in goat over other ruminants and human.

## INTRODUCTION

According to the Global Cancer Report issued by the World Health Organization, there are over 10 million new cases of cancer each year and over 6 million annual deaths from the disease (Chen et al., 2018; Nilendu et al., 2018; Patel et al., 2018; India State-Level Disease Burden Initiative Cancer Collaborators, 2018). Cancer is a type of disease that is caused by an uncontrolled division of abnormal cells in a part of the body. Among various cancer types, breast cancer, cervical cancer, colon cancer and oral cancer are reported as the leading cause of death at global level (Mandel et al., 1993; Nakanishi et al., 2016; Mehta et al., 2017; Nilendu et al., 2017; Bray et al., 2018; Chen et al., 2018; Nilendu et al., 2018; Patel et al., 2018; India State-Level Disease Burden Initiative Cancer Collaborators, 2018). The incidence of these types of cancer in most of countries is projected to rise further over the next 20 years, despite current efforts to prevent the disease for high motility in favor of cancer.

Cancer is commonly treated therapy options such as chemotherapy, surgery, radiation therapy and immune therapy (Mandel et al., 1993; Nakanishi et al., 2016; Mehta et al., 2017; Nilendu et al., 2017; Bray et al., 2018; Chen et al., 2018; Nilendu et al., 2018; Patel et al., 2018; India State-Level Disease Burden Initiative Cancer Collaborators, 2018). These therapies are faced with discernible widely accepted problems including, drug resistance, concomitant its strong side effects such as neurotoxicity, nephrotoxicity, gastrointestinal toxicity vomiting and ototoxicity, recurrence of cancer, poor quality of life post treatment (Mandel et al., 1993; Nakanishi et al., 2016; Mehta et al., 2017; Nilendu et al., 2017; Bray et al., 2018; Chen et al., 2018; Nilendu et al., 2018; Patel et al., 2018; India State-Level Disease Burden Initiative Cancer Collaborators, 2018).

In addition to the above problems, use of these current regimens of drugs is also faced with the social status of patients in developing countries like India where majority of current and future cancer patients are in the weaker economic strata of the society. Hence, viable, sustainable and cheaper source of anti-cancer drugs will be suitable options in Indian and other developing countries.

The role of tumor microenvironment directly and indirectly influenced by microbiotas residing in the local niche of tumor is seeking attention across the globe (Gill et al., 2003; Chino et al., 2010; Stein et al., 2011; Löhr et al., 2013; Sonja et al., 2013; Puthia et al., 2014; Guo et al., 2018; Waluga et al., 2018). It is interesting to know that microbiotas populations differ from individual to individual and also a certain type of microbiotas are known to show anti-cancer and some others are known to act as pro-cancer factors (Gill et al., 2003; Chino et al., 2010; Stein et al., 2011; Löhr et al., 2013; Sonja et al., 2013; Puthia et al., 2014; Guo et al., 2018; Waluga et al., 2018). Approximately, 20% of malignancy are influenced by microbiotas. It is true that this either anti-cancer or pro-cancer group of microbiotas display their effects in tumor microenvironment by releasing several factors including metabolites, peptides, small RNAs and hormones etc.

It is interesting to note that compared to human and other ruminants, goat shows rare clinical evidences of mammary and colorectal cancer (Chino et al., 2010; Stein et al., 2011; Löhr et al., 2013; Sonja et al., 2013; Puthia et al., 2014; Guo et al., 2018; Waluga et al., 2018). The mechanisms behind such observations can be explained by various cellular and molecular phenomena. However, one potential answer can be retrieved in the form of distinct profiles and activities of urine samples from various ruminants and human sources. Additionally, cow urine and goat urine are appreciably mentioned in the Indian traditional medicine (Gill et al., 2003; Chino et al., 2010; Stein et al., 2011; Löhr et al., 2013; Sonja et al., 2013; Puthia et al., 2014; Guo et al., 2018; Waluga et al., 2018).

Therefore, this paper is of the first attempt to understand Xeno-heterogeneity by comparative study on DNA metabolizing and anti-proliferative potential of the DMSO urine fraction of various ruminants including cow, goat, ox, horse, buffalo, jenny and human.

## MATERIAL AND METHODOLOGY

### Materials

Cell culture reagents were purchased from Invitrogen India Pvt. Ltd. and Himedia India Pvt. Ltd. The HCT-116 and MCF-7 cells were procured from National Centre of Cell Science (NCCS), Pune, India. DMSO, pBR322 DNA, agarose, acrylamide and other chemicals were of analytical grade and obtained from Himedia India Pvt. Ltd and Merck India Pvt. Ltd.

### Preparation of UDF from various ruminants and human

Fresh urine samples were collected from healthy ruminates including cow, goat, Buffalo, Ox, Horse Jenny and human in sterile falcon tubes. Then after, fresh urine samples of 20 ml from these sources were subjected to drying at 37°C. After drying of urine samples, sterile DMSO was used to dissolve dried urine samples, pooled together, vortexed and finally centrifuged at 12000Xg to get clear UDF samples in 1 ml DMSO per 20 ml of initial fresh urine samples collected from ruminants and human. The final prepared UDF from ruminants and human was sterile filtered using 0.45 micron syringe to obtain clear and sterile UDF for use in the DNA metabolism activities assay and anti-proliferative assay upon HCT-116 and MCF-7 cells.

### Cell line maintenance and preparation of MCF-7 cells for genomic dna isolation

The HCT-116 and MCF-7 cells were cultured and maintained in DMEM (Dulbecco’s Modified Eagles Medium) (Himedia) with high glucose supplemented with 10% heat inactivated FBS/penicillin (100 units/ml)/streptomycin (100 µg/ml) at 37°C in a humidified 5% CO_2_ incubator. Further, 60-70% confluent MCF-7 cells were harvested and subjected to the genomic DNA isolation as modified protocol (Kumar et al., 2017; Sharma and Kumar, 2018; Sharma et al., 2018). After the isolation of genomic DNA from MCF-7 cells, the integrity and quality of genomic DNA was assessed using standard methods and stored at −20°C for in vitro DNA damaging ability assessment of UDF from various sources.

### In Vitro DNA metabolizing assay

To assess the DNA metabolizing effects of UDF samples from ruminants and human, an in vitro reaction was set up. In this reaction mixture, 1 μl of pBR322 plasmid DNA (100 ng / μl), 1 μg of genomic DNA (MCF-7 breast cancer cells) were mixed with 2.5 μl each of TAE buffer (Tri-acetate/EDTA 10 mM, pH 7.4) and finally treated by different concentration of UDF samples of 1 μl (20 μg to 40 μg per ml) was added to the reaction mixture. The final volume of each reaction mixture was brought to 25 μl by the addition of nuclease free water in a micro centrifuge tube. Reaction mixtures were respectively incubated for 1 h at 37°C. Finally, at the end of incubation, 4 μl of DNA loading dye was added to the incubated mixture, and 20 μl of reaction samples were loaded onto 1% agarose gel. Electrophoresis was conducted at 100 volts in Tris-Acetate-EDTA• Na2 (TAE) buffer (0.04 M Tris-acetate and 1 mM EDTA, pH 7.4) using a Horizon 58 (Life Technologies). At the end of DNA run, agarose gel was stained with ethidium bromide for 15 min and lastly DNA gel was visualized with a Bio-Rad Gel Doc™ EZ imager and densitometry software was used to analyze the DNA metabolized band intensity of plasmid DNA and genomic DNA treated by UDF samples of ruminants and human (Kumar et al., 2017; Jahagirdar et al., 2018; Sharma and Kumar, 2018; Sharma et al., 2018).

### Trypan blue dye exclusion assay

Here, HCT-116 and MCF-7 cells were plated in six well plate (1.5*105 cells per well) and incubated in the presence of complete fresh DMEM medium supplemented with 10% FBS. After 16-18 h, cells were treated with 2 ml of complete fresh DMEM medium and added with varied concentration (10, 50 and 100 µl/ml) final concentration of goat UDF sample (stock 10 mg per ml). (10, 50 and 100 µl/ml) final concentration. A negative control was also prepared by including complete fresh DMEM medium mixed with equal amount of DMSO solvent. After 72 h of incubation, the media was aspirated to recover the floating cells. Next, cells were washed with PBS and treated with 0.25% trypsin-EDTA for 2-3 min at 37ºC. Fresh media were added into each well and cells were harvested properly. To check the total and viable cell count in the collected cell suspensions, a routinely used Trypan blue dye exclusion was performed to estimate viable cell and dead cell using Hemocytometer.

### STATISTICAL ANALYSIS

Data shown are presented as the mean ± SD of at least three independent experiments; differences are considered statistically significant at P < 0.05, using a Student’s t-test.

## RESULTS AND DISCUSSION

### DNA metabolizing effects of UDF from various sources

In this paper, UDF samples from various ruminants and human are prepared and filtered in a sterile condition. Based on the literature, distinct intestinal microbiotas and potential diet derived metabolites are known to be distinct from each other. However, a simple assay to know the DNA metabolizing effects of these UDF composition is not reported and this assay may provide additional evidence in support of distinct of UDF samples among all these sources of urine samples.

To assess the DNA metabolizing effects of UDF samples, Figure 1A shows a photograph of agarose gel electrophoresis-based separation of in vitro plasmid DNA pBR322 that is treated with UDF samples from various ruminants and human. The DNA band intensity of plasmid DNA after the treatment by UDF from various ruminants and human is presented as a bar graph in Fig, 1B. In this result, DMSO serves as solvent control and data suggest the presence of very clear DNA metabolizing effects up to 92% by goat UDF and found to be maximum compared over other ruminant and human UDF samples. Here, authors do not claim the direct distinctiveness of chemical composition of goat UDF over others. But this DNA metabolizing assay strongly indicates the possible nature of unique compositions possibly as metabolites secreted in the goat UDF over other UDF samples. Further, it is important to mention that metabolized DNA damage pattern by goat UDF is distinct from one of other cow UDF samples that displayed lesser extent of DNA metabolizing effects. Interestingly, authors suggest that DNA metabolizing effects observed in a particular reaction buffer should not be seen as a source of carcinogenic substances or heavy metal contaminations. We conducted other supporting experiments and data indicate that DNA metabolizing effects are only observed in a particular buffer mentioned in this paper. Besides physiological buffers used in cell culture, we do not see discernible DNA damaging effects as seen in TAE buffer condition (data not shown). From these experiments, we indirectly infer that goat UDF has distinctive chemical compositions that become active in terms of their functional groups and possibly demonstrate very strong DNA metabolizing effects.

**Fig. 1.**
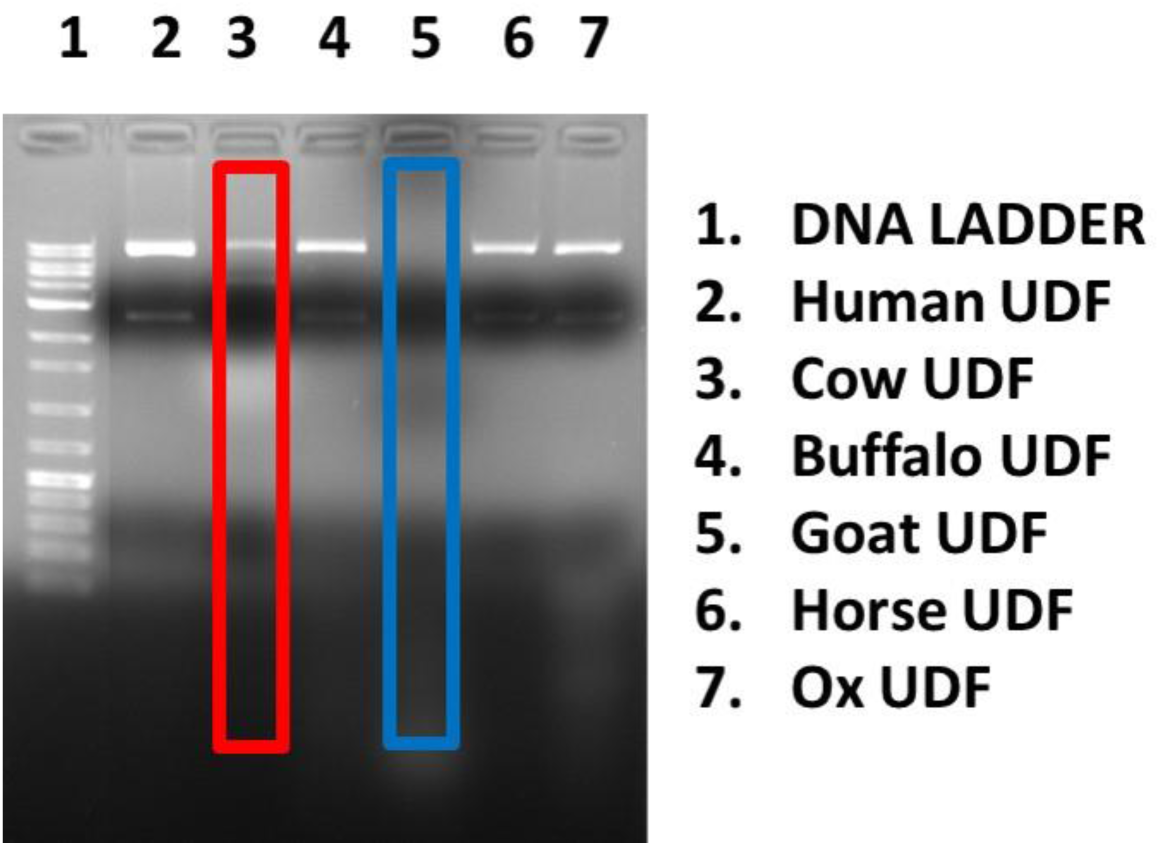
DNA metabolism effects of UDF samples from various sources upon pBR322 plasmid DNA. This photograph shows an ethidium bromide stained agarose gel electrophoresis-based separation of in vitro plasmid DNA pBR322 treated UDF from various ruminants including cow, goat, buffalo, ox, horse and also human. The image was visualized and captured using BIO-RAD EZ imaging system. Fig.2B show the bar graphs as percentage of analyzed pBR322 plasmid DNA in terms of total circular DNA including closed circular and relaxed plasmid DNA over DMSO control. The bar graph without any asterisk denotes that there is no any significant difference compared to DMSO control. * Significantly different from DMSO control at P-value < 0.05.

Similarly, Figure 2A represents a photograph of agarose gel electrophoresis-based separation of genomic DNA derived from MCF-7 breast cancer cells that is treated by UDF samples from various sources. The genomic DNA instability is analyzed using the band intensity densitometry analysis software and data is given as a bar graph in Fig. 2B. The analysis of data in Fig. 2B indicates highly appreciable genomic DNA degradation in the range of 80 to 90% goat UDF compared to UDF samples from other ruminants and human. Among many ruminants and human samples, cow UDF sample display to some extent of DNA metabolizing effects upon the genomic DNA, however, the pattern of DNA damage to genomic DNA is clearly distinct from goat UDF to cow UDF. In this data, it is mentioned that the amount of used UDF samples ranged from 20 to 40 µg per ml and the volume of UDF samples used in this reaction ranged from 0.5 µl to 1 µl per reaction volume. The observed differential DNA metabolizing effects of goat and cow UDF in comparison to other ruminants and human UDF samples is well explained by the presence of distinct metabolites in the isolated UDF samples from various ruminants and human. Importantly, DNA metabolizing effects of precisely goat and cow UDF can be attributed to majorly to metabolites and similar compounds as protein profile details do not support the presence of proteins and urea and this may be due to the nature of fractionation process and solvent DMSO that exclude the possibilities of urea, similar to urea and protein components.

**Fig. 2.**
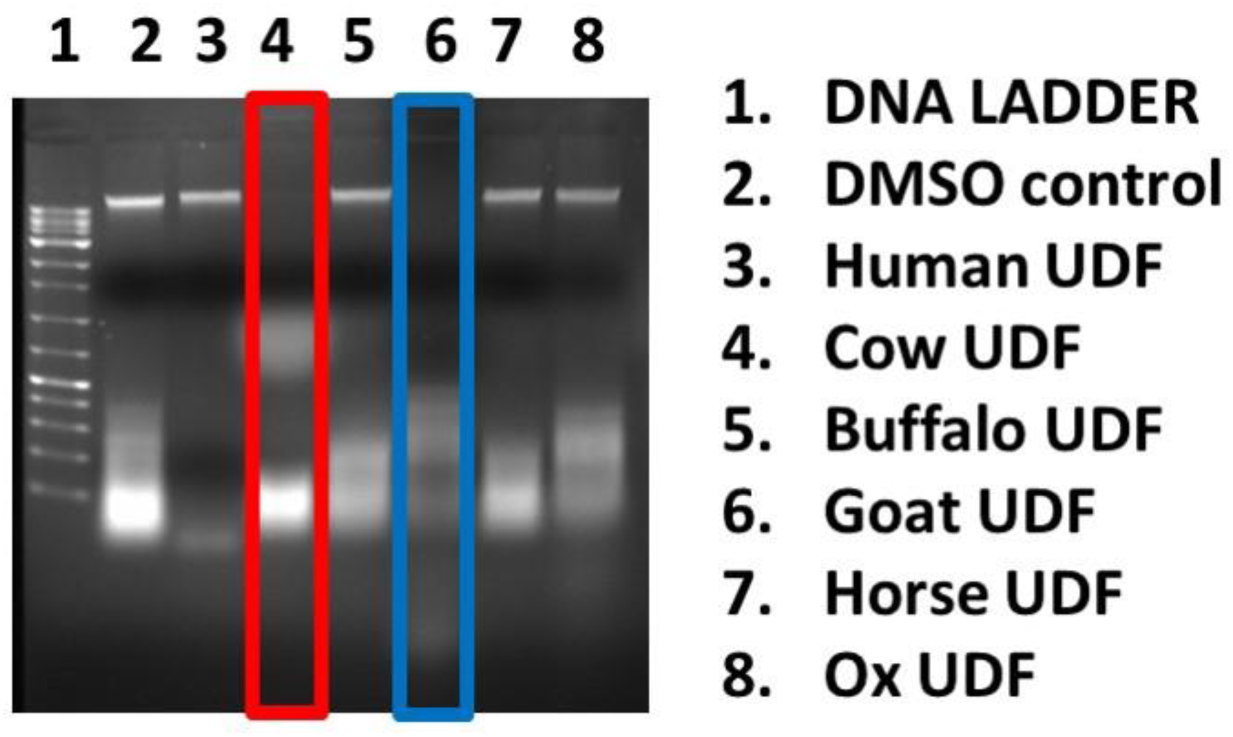
DNA metabolism effects of UDF samples from various sources upon genomic DNA from MCF-7 breast cancer cells. In Fig. 2A. photograph depicts an ethidium bromide stained agarose gel electrophoresis-based separation of in vitro genomic DNA from MCF-7 breast cancer cells treated by UDF from various ruminants including cow, goat, buffalo, ox, horse, Jenny and also human. The image was visualized and captured using BIO-RAD EZ imaging system. Fig. 2B show the bar graphs as percentage of analyzed genomic DNA band intensity over DMSO control and various UDF samples from ruminants and human. The bar graph without any asterisk denotes that there is no any significant difference compared to DMSO control. * Significantly different from DMSO control at P-value < 0.05.

In this reference, a concern is raised that goat UDF sample may contain some inorganic contaminations including the presence of heavy metal ions. To address and exclude such possibilities, autoclaved UDF samples are subjected to similar DNA metabolizing assay. Interestingly, data from Fig. 3A and Fig. 3B clearly indicate the almost absence of DNA metabolizing effects after the autoclaving of goat UDF sample and other samples. In this study, both autoclaved and non-autoclaved UDF samples were filtered by using sterile 0.45 µm size membrane. Surprisingly, data clearly support that the nature of DNA metabolizing components from goat UDF and other samples exclude the possibilities of heavy and other inorganic compounds contaminations At the same, an indirect evidence to the presence of metabolites such organic acids, short chain fatty acids and other kinds of metabolites. Further, LC-MS and other suitable analytical assays are in progress to precisely identify the chemical compositions in the goat UDF sample.

**Fig. 3.**
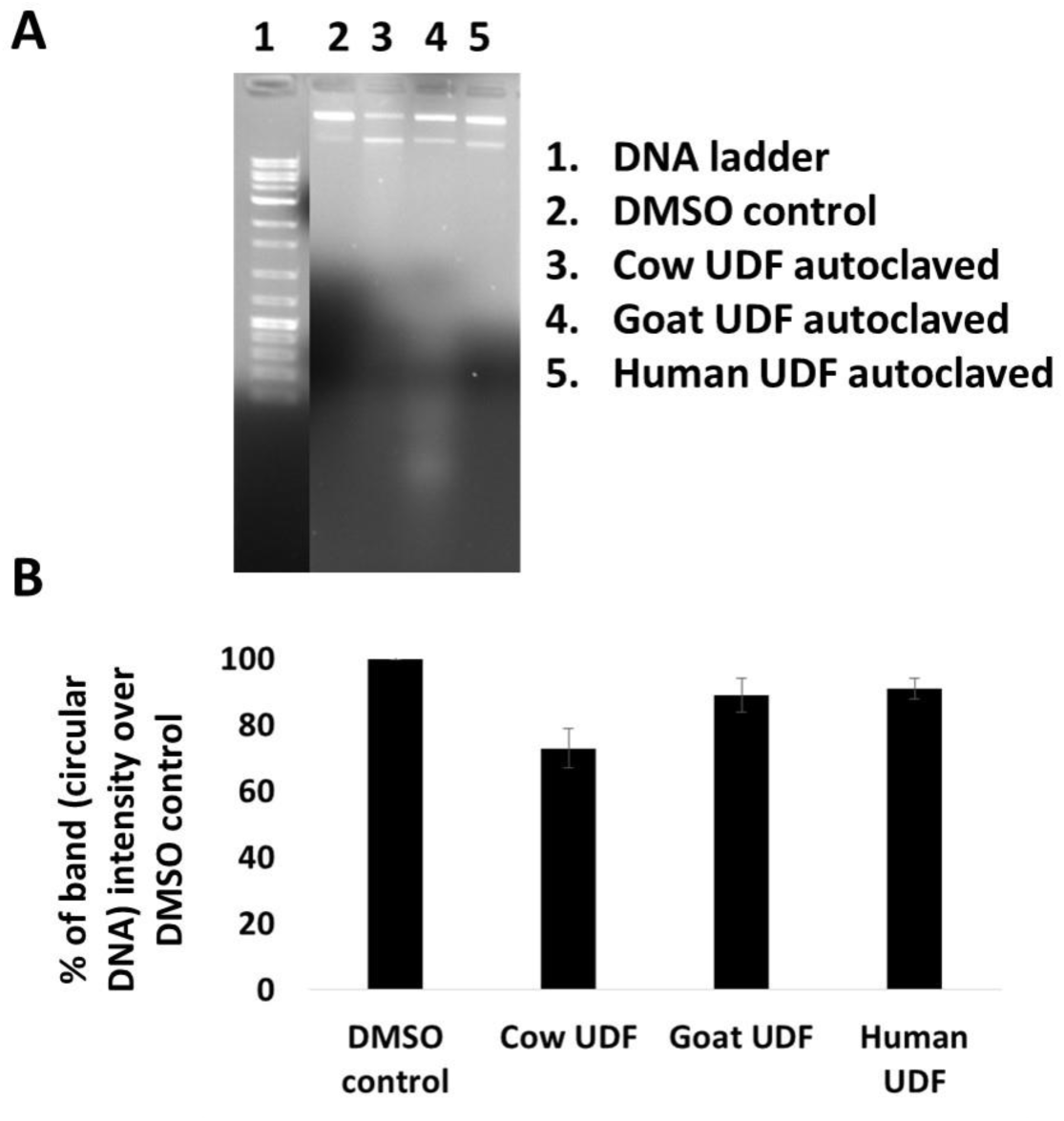
Loss of DNA instability effects by autoclaved UDF samples from cow, goat and human Fig. 3A photograph illustrates an ethidium bromide stained agarose gel electrophoresis-based separation of in vitro plasmid DNA pBR322 treated by autoclaved UDF from cow, goat, and human. The image was visualized and captured using BIO-RAD EZ imaging system. Fig.3B show the bar graphs as percentage of analyzed pBR322 plasmid DNA in terms of total circular DNA including closed circular and relaxed plasmid DNA over DMSO control. The bar graph without any asterisk denotes that there is no any significant difference compared to DMSO control. * Significantly different from DMSO control at P-value < 0.05.

Based on the very clear DNA metabolizing effect of goat UDF and also the literature that suggest that goat is known to show the high resistance to colorectal and mammary cancer and rare occurrence of these cancer types (Chino et al., 2010; Stein et al., 2011; Löhr et al., 2013; Sonja et al., 2013; Puthia et al., 2014; Guo et al., 2018; Waluga et al., 2018). The authors indicate that the DNA metabolizing effect of goat UDF is potentially correlated and may serve as an indicator of distinctive metabolites or organic acid components in goat UDF sample. Therefore, we evaluated anti-proliferative effect of goat UDF upon HCT-116 and MCF-7 cancer cells. This anti-proliferative effect of goat UDF is assessed by using Trypan blue due exclusion assay, which is a routine, reproducible and simple experiment. In essence, data from Trypan blue dye exclusion method represent the status on total cell count and percentage of cell survival induced by the incubation of growing HCT-116 and MCF-7 cells in the presence of goat UDF. Here, the authors present microscopy photographs in both normal contrast and phase contrasts (Figure 4A).

**Fig 4.**
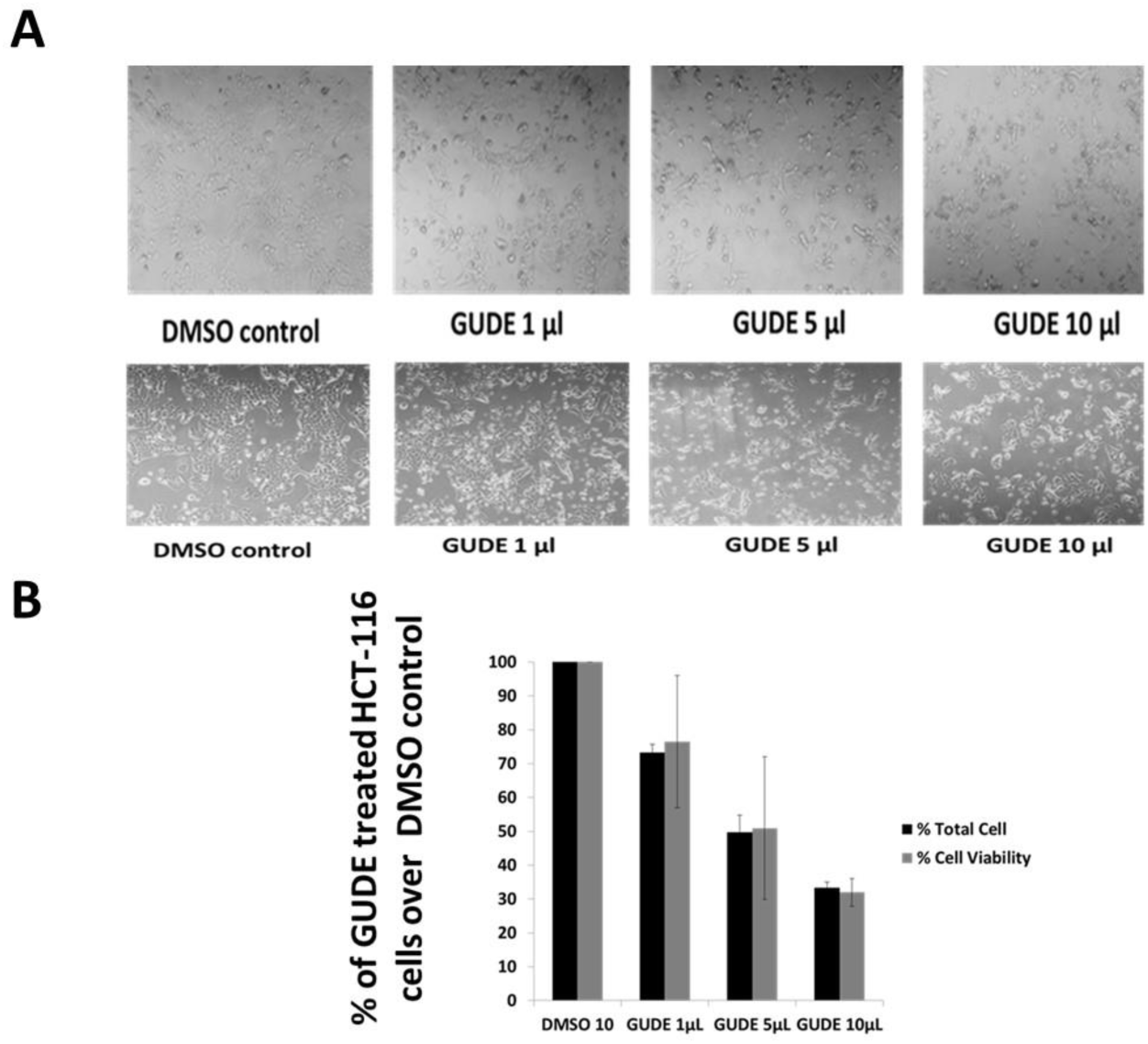
(A) Microscopic photographs taken at 100X at the end of 72hr of treatment to HCT-116 cells growing in six well plate with varied concentration (10, 50 and 100 µg/ml) final concentration of goat UDF sample (stock 10 mg per ml). (4B) Bar graph data represents the total cell and viable HCT-116 cell count at the end of 72 hr of treatment with goat UDF sample. The data are represented as mean ± SD. Each experiment was conducted independently three times.

The analysis of photomicrographs convincingly indicates the significant decrease in the total number of HCT-116 cells and also the changes in the morphology of treated HCT-116 cells. Further, a graphical representation of total HCT-116 cells and total viable HCT-116 cells is shown in Fig. 4B. The data clearly shows the highly significant decrease in the total number of HCT-cells and also the total number of viable HCT-116 cells. In essence, this cell based assay does not provides the information on the nature and type of cell death due to treatment by goat UDF. As earlier suggested by several other reports, Trypan blue dye exclusion assay provides information on anti-proliferative and anti-survival potentials, but may not be sufficient to reveal the kind of cell death and other cellular changes including cell cycle changes in treating cancer cells. In this paper, current preliminary data appears to be sound and reproducible to support the effects of goat UDF sample upon HCT-116 cells. Next, we tested the anti-proliferative effects upon other cell line MCF-7 cells, which is completely distinct from the HCT-116 cells in terms of genetic and molecular adaptations. In this way, the authors are interested to know that whether distinctive composition of goat UDF will display anti-proliferative effects upon any types of cancer cells. Surprisingly, data collected from using Trypan blue dye exclusion assay upon MCF-7 cells suggest that anti-proliferative efficacy of goat UDF is significantly less compared to HCT-116 cells (Fig. 5A and Fig. B).

**Fig 5.**
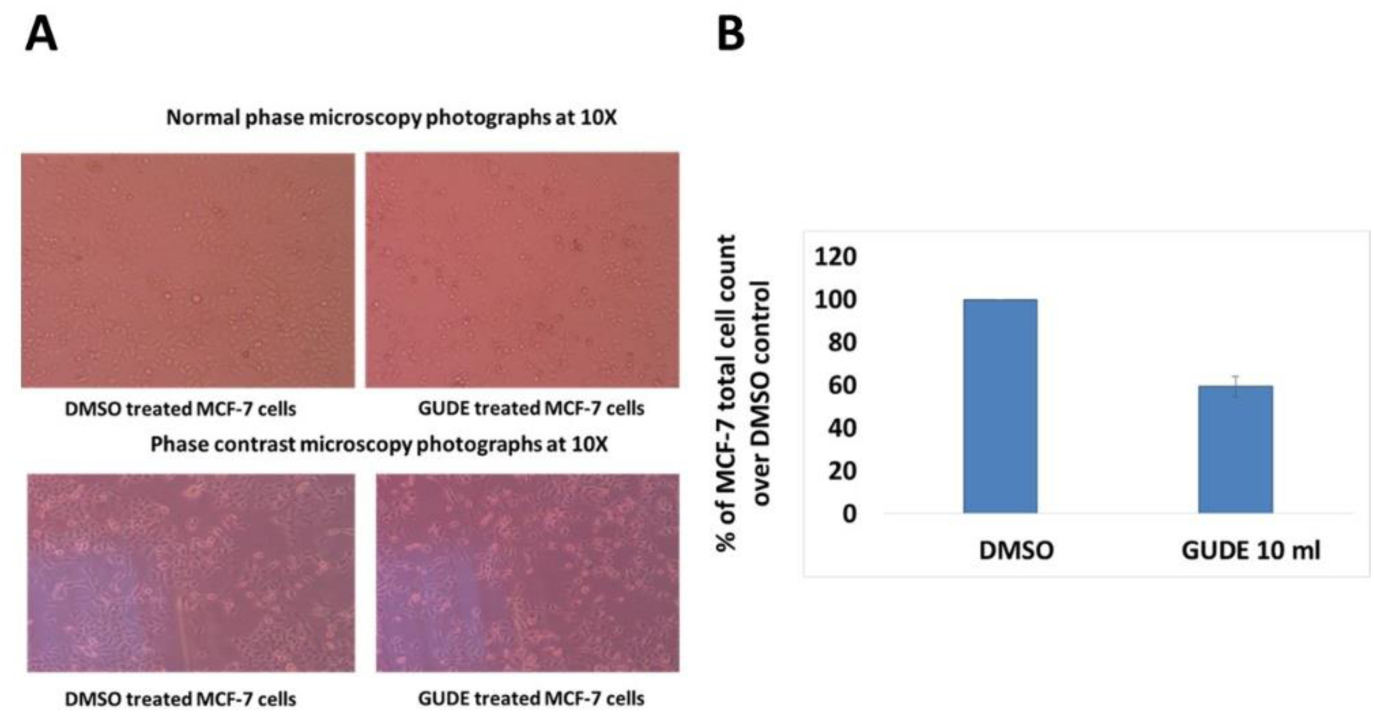
(A) Microscopic photographs taken at 100X at the end of 72hr of treatment to MCF-7 cells growing in six well plate with final concentration (100 µg/ml) of goat UDF sample (stock 10 mg per ml). (5B) Bar graph data represents the total cell and viable of MCF-7 cell count at the end of 72 hr of treatment with goat UDF sample. The data are represented as mean ± SD. Each experiment was conducted independently three times.

At this juncture of preliminary observations, we speculate that the distinctive goat UDF composition may have specific targets like nuclear receptor in the HCT-116 cells compared to MCF-7 cells, which may be able to contribute to better anti-proliferative effects in MCT-116. Interestingly, rare occurrence of colorectal cancer in goat, unique microbial colonization of goat gut system, and derived metabolites in goat urine converge upon to indicate that the observed findings from in vitro anti-proliferative effects of goat UDF compared to UDF samples from other ruminants and human may have preclinical and clinical relevance. On the future road map, these goat UDF and other samples are in process of LC-MS analysis to confirm the chemical identities that may be a contributing factor in the observed findings in DNA metabolizing and anti-proliferative effects upon HCT-116 cells. Further, this finding may set the platform to investigate on goat UDF samples as potential novel potential anti-cancer pharmaceutical composition. The possible claim of this paper is well supported from the basic understanding of cancer biology that depicts cancer is highly complex and heterogeneous nature. In recent, tumor heterogeneity and various hallmarks such as drug resistance is well explained by the distinct heterogeneity at cellular levels that include various types of cells like immune cells, stromal cells, etc. Besides these known cells, presence of microbiotas in the local niche of tumor is well associated to the tumor hallmarks and tumor progression. Interestingly, different populations of microbiotas are reported to act as both tumor promoters and tumor inhibitors.

In a comprehensive review, various ruminants and human individuals are evidenced to display distinct gut microbiotas and related secreted metabolite profiles in urine samples from these sources. Our finding also supports that may be distinct gut microbiotas in various ruminants and as well distinct nutrition factors contribute towards differential urine metabolites and these urine metabolites show distinct activities as reported in this paper as DNA metabolizing effects of cow and goat UDF.

### FUTURE DIRECTIONS

In future, studies can be conducted to validate the in vitro effects of goat UDF in colon cancer in mice model. In future, metabolites components of goat UDF can be combined with nano-carriers for better drug delivery. In a better quest, molecular targets like nuclear receptors of colorectal cancer cells may be explored as specific targets of metabolites components of goat UDF and further warrants study in both in vitro and in vivo colorectal cancer model.

## SIGNIFICANCE STATEMENT

In this paper, the authors draw the attention to suggest that this simple finding is potentially linked to one of key question tumor heterogeneity in cancer biology. Here, findings propose that goat and other ruminants may have naturally equipped metabolic systems contributed by gut system, microbiotas in the local niche and dietary derived metabolites to thwart carcinogenesis process in goat. Conversely, human and other highly susceptible vertebrates may lack these anti-proliferative and anti-cancer metabolites derived from gut system, microbiotas in the local niche and dietary derived metabolites. In essence, human and other highly susceptible vertebrates may generate pro-cancer and pro-carcinogenesis metabolites from gut system, microbiotas in the local niche and dietary derived metabolites. Taken together, this findings warrants other preclinical scientists to look into detailed mechanisms that may lead to the better understanding on tumor heterogeneity, revealing novel metabolite biomarkers and therapeutic interventions in future.

## CONCLUSION

In conclusion, goat UDF displays distinct abilities to metabolize plasmid and genomic DNA substrates. Further, goat UDF treatment to HCT-116 shows very strong anti-proliferative effects and warrants appreciable future investigations in vitro and in vivo model of colorectal cancer. Interestingly, goat UDF does bring similar extent of effects upon MCF-cells. It is important to conclude that goat UDF sample used for the present DNA metabolizing and anti-proliferative assay is filter membrane sterilized. But, autoclaving of goat UDF sample shows the loss of both DNA metabolizing and anti-proliferative effects. This finding supports the one of the possible answer behind the xeno-tumor heterogeneity among ruminants and human idea that may establish definite connections with differential occurrence (rare in goat) and high occurrence (human).

## Acknowledgements

The authors acknowledge financial support from DST-SERB, Government of India, New Delhi, India (SERB/LS-1028/2013) and Dr. D. Y Patil, Vidyapeeth, Pune, India (DPU/05/01/2016). The authors acknowledge Central Research Facility, DPU, Pune for flow cytometer and fluorescent microscopy experimentation facilities.

